# TMEM100, a Lung-Specific Endothelium Gene

**DOI:** 10.1101/2022.08.26.504609

**Authors:** Bin Liu, Dan Yi, Zhiyun Yu, Jiakai Pan, Karina Ramirez, Shuai Li, Ting Wang, Christopher C. Glembotski, Michael B. Fallon, S. Paul Oh, Mingxia Gu, Joanna Kalucka, Zhiyu Dai

## Abstract

The heterogeneity of endothelium across different organs was recently explored using single-cell RNA-sequencing analysis. Compared to other organs, the lung exhibits a distinct structure composed of a thin layer of capillary for efficient gas exchange. In this study, we demonstrate that Tmem100 is a lung-specific endothelium gene.

---

Endothelial cells (ECs) line the inner wall of blood vessels and maintain tissue homeostasis by preserving vascular integrity via blood flow regulation and exchanging oxygen and nutrients. The heterogeneity of ECs across different organs was recently explored using single-cell RNA-sequencing (scRNA-seq) by multiple studies^1^. Compared to other organs, the lung exhibits a distinct morphology composed of a thin layer of capillary ECs for efficient gas exchange. Despite their vital role in lung function, neither a unique lung-specific EC gene nor a lung-EC specific Cre line has been identified in the literature. In this study, we demonstrate that Tmem100 is a lung-specific endothelium gene using scRNA-seq and lineage tracing approaches.

Via leveraging the recent murine ECs scRNA-seq dataset^1^, we found that *Tmem100* is one of the top marker genes distinguishing lung ECs from other organs ECs such as heart, brain, kidney, small intestine, etc (Figure [A], i). We further extracted ECs transcriptomes from Tabula Muris scRNA-seq dataset^2^, and confirmed that *Tmem100* is the lung EC-specific gene (Figure [A], ii). The expression of Tmem100 and VE-cadherin protein was validated across different organs by Western Blotting (Figure [A], iii). General ECs are characterized by uniform expression of vascular endothelial (VE)-cadherin (*CDH5*). To determine whether *TMEM100* is selectively expressed in the human lung ECs compared with other cell types in the lung, we compared the expression pattern of *TMEM100* and *CDH5* in the scRNA-seq data from 12 healthy human lungs^3^. The expression pattern of *TMEM100* is in parallel with *CDH5* (Figure [A], iv), suggesting that *TMEM100* is selectively expressed in lung ECs. Afterwards, we de novo analyzed lung scRNA datasets generated from adult mouse and rat (including both male and female) lungs in house and confirmed that *Tmem100* is an EC selective marker in the rodent lung (data not shown).

**Figure.**
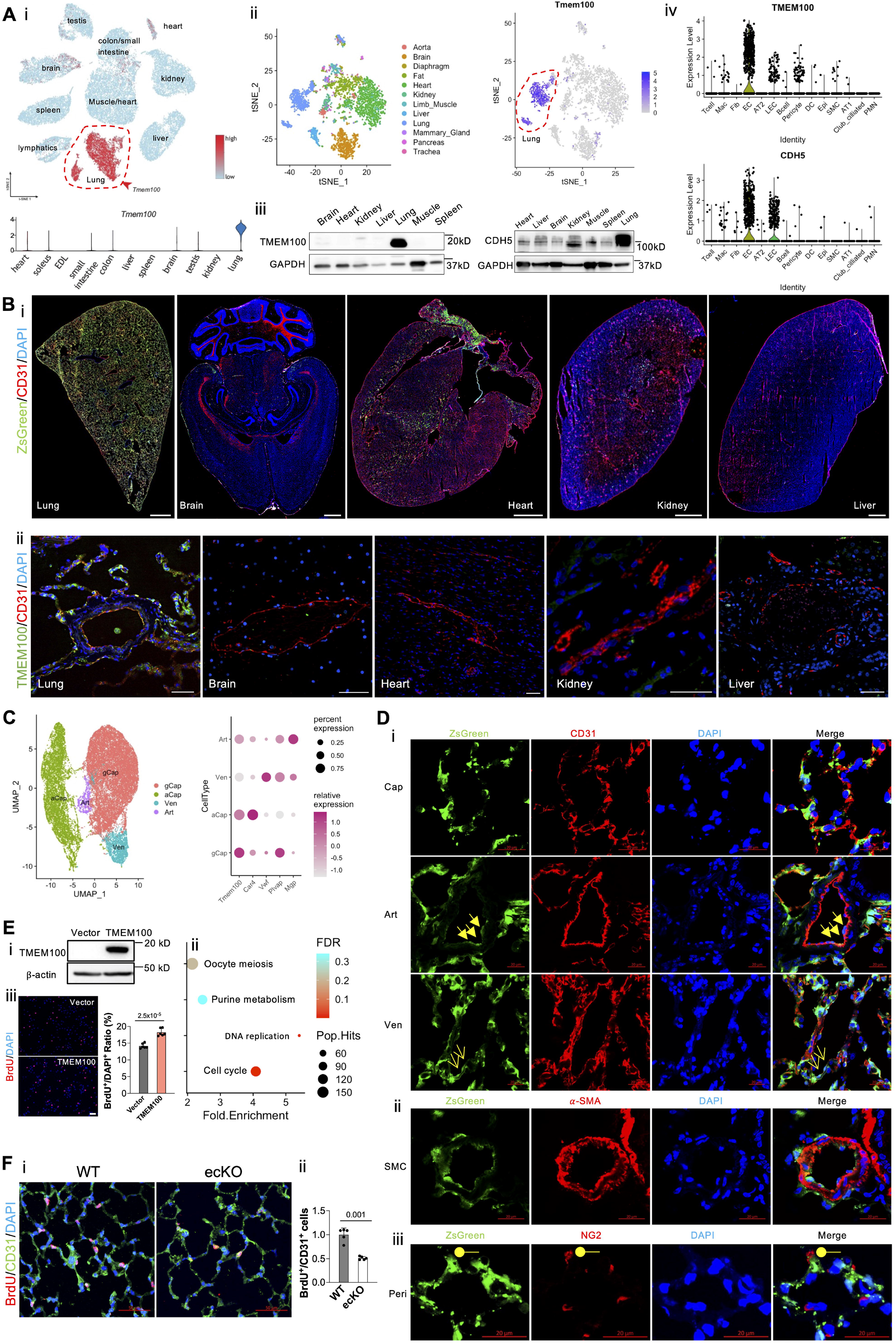
Tmem100, a lung-specific endothelium gene. **(A**, i). A representative tSNE plot and a violin plot of murine ECs scRNA-seq data showing the lung EC-specific expression of *Tmem100* in comparison to other organs’ ECs. (**A**, ii**). A** representative tSNE plot of extracted EC transcriptomic analysis of Tabula Muris scRNA-seq dataset demonstrating unique *Tmem100* expression pattern in lung ECs. **(A**, iii). Western Blotting demonstrated Tmem100 protein selectively expressed in the mice lung. GADPH was used as a loading control. **(A**, iv). A representative Violin Plot showing that *TMEM100* was selectively expressed in the lung ECs similar to classical EC marker *CDH5* in human. **(B**, i). A lineage tracing strategy using *Tmem100*^*ZsG*^ mice demonstrated that Tmem100 (ZsGreen^+^) was enriched in the lung ECs (CD31^+^) but not the ECs in other organs such as brain, heart, kidney, and liver. Both male and female *Tmem100*^*ZsG*^ mice at the age of 7 weeks were injected intraperitoneally with tamoxifen daily for 5 days (20mg/kg, daily) and rested for ∼10 days. Scale bar: 1 mm. **(B**, ii). Immunostaining demonstrated TMEM100 is selectively expressed in human lung ECs. Scale bar: 50 *μ*m **(C)**. A UMAP plot showing the different subtypes of lung ECs based on the murine EC dataset, and *Tmem100* highly expressed in general capillary EC (gCap), aerocytes (aCap), and lowly expresses in arterial (Art) and venous (Ven) ECs. (**D**, i) Immunostaining demonstrated that ZsGreen^+^ (Tmem100^+^) was strongly expressed in the capillary ECs and weakly expressed in the arterial and venous ECs. Cap, capillary EC; Art, arterial EC (triangle arrows); Ven, venous EC (arrows). (**D**, ii) ZsGreen^+^ (Tmem100^+^) was not expressed in SMCs (*α*-SMA^+^). (**D**, iii) A few NG2^+^ cells (pericytes, Peri, round shape) expressed ZsGreen (Tmem100). Scale bar: 20 *μ*m. **(E)**. TMEM100 overexpression (**E**, i) in HPMVECs induced enrichment of pathways related to Cell Cycle, DNA Replication (**E**, ii) based on the upregulated genes induced by TMEM100 overexpression. Three replicates were pooled in the equal amount for RNA-seq analysis. (**E**, iii). TMEM100 overexpression promoted EC proliferation accessed by BrdU assay. At 48 hours post-lentiviruses infection, HPMVECs were starved in serum/growth factors free medium for 12 hours. BrdU (10*μ*M) was added in the medium at 4 hours prior to cells harvest. BrdU was stained with anti-BrdU antibodies. Scale bar: 100 *μ*m. N=6 (biological replicates). Student t test. (**F**, i, ii). Genetical deletion of Tmem100 in mice impaired EC proliferation in vivo during inflammatory lung injury. *Tmem100*^*f/f*^ (WT) and Tmem100^EndoSCL-CreERT2^ (ecKO) mice were treated with LPS (5mg/kg) intraperitoneally. Mice were injected BrdU (25 mg/kg) intraperitoneally daily for two days. Lung tissue were collected at 72 hours post-LPS treatment. BrdU was stained with anti-BrdU antibodies. N=5 (biological replicates). Welch’s t-test. Statistics. (GraphPad Prism 9.0): based on additional literature support from similar studies, our samples fit normal distribution. Parametric test: parametric test with the unpaired 2-tailed Student t test for equal variance and Welch’s t-test for unequal variance (2 groups). Mean± SD.

To further determine the EC-selective expression of *Tmem100 in vivo*, we generated *Tmem100*^*ZsG*^ mice via breeding *Tmem100-CreER*^*T2*^ mice (JAX: 014159) with *ZsGreen*^*LSL/LSL*^ mice (JAX: 007906). The expression pattern of ZsGreen fluorescence indicates Tmem100 expression in mice after tamoxifen (20mg/kg, 5 doses, daily) treatment. We then visualized ZsGreen fluorescence together with pan EC marker CD31 in multiple organs including lung, heart, brain, kidney, and liver cryosections. Full organ slides imaging of ZsGreen showed that ZsGreen expression is robustly expressed in the lung and the right atrium compared to other organs examined (Figure [B], i). We also determine the expression of TMEM100 in different human organs and confirming that *Tmem100* is predominantly a gene expressed in lung ECs (Figure [B], ii).

Following examination of the expression pattern of ZsGreen in the lung, we found that the ZsGreen^+^ cells colocalized with 96% CD31^+^ cells in the lung quantified by flow cytometry (data not shown). Recent studies employing scRNA-seq analysis demonstrated that lung ECs contain arterial (expressing markers *Mgp, Sparcl1*)^1^, venous (*Prss23, Vwf*)^1^, and general capillary ECs (*Gpihbp1, Plvap*), and aerocytes (*Car4, Ednrb*)^1,4^. Using similar markers to define EC subpopulations, our re-analyzed scRNA data showed that *Tmem100* is higher in general capillary EC and aerocytes, and lower in arterial EC and venous ECs (Figure [C]). Immunostaining and quantification analysis confirmed that ZsGreen^+^ cells are robustly expressed in the capillary ECs (92%) and weakly expressed in the arterial (72%) and venous ECs (48.6%). (Figure [D], i). Human lung scRNA-seq data (Figure [A], iv) suggested *TMEM100* was also expressed in some pericytes and smooth muscle cells (SMCs). We then stained our lung tissue with the pericyte marker NG2 and SMC marker *α* -SMA and found that not *α* -SMA^+^ cells (Figure [D], ii), but a few NG2^+^ cells (1%) (Figure [D], iii) colocalized with ZsGreen^+^ cells.

Fresh isolated mouse ECs express high TMEM100, whereas human pulmonary arterial ECs (PAECs) or microvascular ECs (PMVECs) do not express detectable TMEM100 in 2-dimension culture by Western Blotting. To overcome this limitation, we overexpressed TMEM100 fused with 3XFlag tag (synthesized by Genscript) via lentivirus (vector from Addgene:36084) in HPMVECs for RNA-seq analysis (Figure [E], i). We then performed enriched pathway analysis via Kyoto Encyclopedia of Genes and Genomes (KEGG) on the upregulated genes by TMEM100 overexpression, which pointed to Cell Cycle and DNA replication pathways (Figure [E], ii). TMEM100 overexpression promoted HPMVECs proliferation in vitro. (Figure [E], iii). Genetic deletion of Tmem100 in ECs impaired EC regeneration after endotoxin challenge (LPS, 5 mg/kg) in mice. (Figure [F], i, and [F], ii). This observation is consistent with a previous finding that *Tmem100* deficiency in mice reduced retinal angiogenesis^5^. We did not observe EC junction integrity change by TMEM100 overexpression by accessing trans-endothelial resistance *in vitro*.

Via incorporating multiple scRNA-seq data from human and rodents, and lineage tracing analysis, our data proposed that *Tmem100* as a selective lung EC marker. In addition to the existing EC-specific labeling Cre lines, *Tmem100-CreER*^*T2*^ mice might be an ideal tool for dissecting the role of lung specific ECs without affecting vascular beds in the organs outside the lung.

## Acknowledgements

All animal protocols were approved by the Institutional Animal Care and Use Committee of University of Arizona. This study used samples, data, and/or services from the Discover Together Biobank at Cincinnati Children’s Research Foundation. We thank the Discover Together Biobank for support of this study, as well as participants and their families, whose help and participation made this work possible. The authors thank the Pulmonary Hypertension Breakthrough Initiative (PHBI) for providing the lung tissues. Funding for the PHBI is provided under an NHLBI R24 grant (R24HL123767). Raw data and complete methods are available upon request from the corresponding author. RNA-seq data have been deposited in the GEO database under accession number GSE209868. Data not shown in the main figure can be accessed via: https://www.biorxiv.org/content/10.1101/2022.08.26.504609v1.

## Highlights

- **Tmem100 is enriched in lung endothelial cells**
- **Tmem100-CreER**^**T2**^ **mice model is an ideal tool to label lung endothelial cells**
- **Lung endothelial Tmem100 is required for endothelial regeneration**

