## supplemental data for "TMEM100, a Lung-Specific Endothelium Gene"

### Online Methods

#### Data availability

RNA-seq data have been deposited in the GEO database under accession number GSE209868. Scripts used for single-cell RNA sequencing analysis and analyzed data in R objects are available in Figshare (https://doi.org/10.6084/m9.figshare.20430753.v1). Other data that support the findings of this study are available from the corresponding author upon reasonable request.

#### Animals

*Tmem100-CreER*$^{T2}$ mice (JAX: 014159) and *ZsGreen*$^{LSL/LSL}$ mice (JAX: 007906) were purchased from JAX lab. *Tmem100*$^{ZsG}$ (*Tmem100-CreER*$^{T2}$*; ZsGreen*$^{LSL/+}$) mice were generated via breeding *Tmem100-CreER*$^{T2}$ mice with *ZsGreen*$^{LSL/LSL}$ mice. We did not observe any difference between male and female mice regarding Tmem100 expression based on the Tabula Muris EC scRNA-seq analysis (**Supplemental Figure F**). To minimize the use of animals, we did not separate male and female mice. Both male and female *Tmem100*$^{ZsG}$ mice at the age of ~7 weeks were treated with Tamoxifen (20mg/kg, daily) in corn oil intraperitonially for 5 days. The lung tissues were collected at 10 to 15 days post final Tamoxifen treatment.

*Tmem100* floxed mice were gifted by Dr. S. Paul Oh (Barrow Neurological Institute). *Tmem100*$^{EndoCreERT2}$ mice (ec KO) were generated by crossing *Tmem100* floxed mice with EndoSCL-CreER$^{T2}$ transgenic mice expressing the tamoxifen- inducible Cre recombinase under the control of the 5′ endothelial enhancer of the stem cell leukemia locus. For endotoxin challenge induced acute lung injury, *Tmem100*$^{f/f}$ (WT) and ecKO mice were treated with Tamoxifen to induce Cre expression and Tmem100 deletion. Three weeks post tamoxifen treatment, both male and female WT and ecKO mice were injected with LPS (Sigma, Cat# L2630-10MG, 5mg/kg) intraperitoneally. To assess in vivo proliferation, mice were injected BrdU (75 mg/kg) intraperitoneally daily for two days. Lung tissue were collected at 72 hours post-LPS treatment. The animal care and study protocols were reviewed and approved by the Institutional Animal Care and Use Committees of the University of Arizona.

#### Human samples

The use of human samples was approved by the University of Arizona Institutional Review Board and Cincinnati Children's Medical Center. Human tissues used in this study were obtained from the Discover Together Biobank at Cincinnati Children's Research Foundation and Pulmonary Hypertension Breakthrough Initiative (PHBI). The sample information including age and sex were provided in Supplemental Table 1.

#### Reanalysis of public single-cell RNA sequencing datasets

We used the publicly available single-cell RNA sequencing metadata from human lung (GSM3926539)[1]. The metadata was processed in R (version 4.0.2) via Seurat package V4 [ref]. Briefly, cells that expressed fewer than 100 genes, and cells that expressed over 4,000 genes, and cells with unique molecular identifiers (UMIs) more than 10% from the mitochondrial genome were filtered out. The data were normalized and integrated in Seurat, followed by Scaled and summarized by principal component analysis (PCA), and then visualized using Uniform Manifold Approximation and Projection (UMAP) plot. FindClusters function (resolution = 0.3) in Seurat was used to cluster cells based on the gene expression profile. Endothelial cells (expressing CDH5, PECAM1), fibroblasts (FN1, COL1A1), smooth muscle cells (ACTA2, MYH11), type 1 alveolar epithelial cells (HOPX, AQP5, AGER), type 2 alveolar epithelial cells (SFTPC), club ciliated cells (SCGB1A1), pericytes (PDGFRB), Mast cells (MCTP1), PMN (S100A9, S100A8) and macrophages (CD68, CD14), T cells (CD3G), dendric cells (IRF8, MS4FA1) clusters were annotated based on the expression of known markers. We also re-analyzed the mouse EC single-cell RNA sequencing dataset[2] and Tabula Muris scRNA-seq dataset[3,4].

**Immunostaining**

Mouse lung tissues were perfused with PBS, inflated with 50% OCT in PBS, and embedded in OCT for cryosectioning. Both human and mouse lung sections (5 μm) were fixed with 4% paraformaldehyde and blocked with 0.1% Triton X-100 and 5% normal goat serum at room temperature for 1 hour. After 3 washes with PBS, the slides were incubated with anti-TMEM100 (Millipore Sigma, Cat#ABN1721, 20 $\mu$g/ml) or anti-CD31 (R&D, Cat#AF806, 10$\mu$g/ml, for human slides), anti-CD31 antibody (BD Bioscience, Cat#550274, 0.625 $\mu$g/ml, for mouse slides), anti-α-SMA (Abcam, Cat#Ab5694, 3.3 $\mu$g/ml), anti-NG2 (Santa Cruz, Cat#SC-166251, 8 $\mu$g/ml) at 4°C overnight then incubated with Alexa 594 conjugated secondary antibodies at room temperature for 1 h. Nuclei were counterstained with DAPI contained in mounting media.

All the antibodies used in this study were carefully selected and validated. The antibodies used in our studies including anti-BrdU, anti-CD31, anti-α-SMA were previously validated by immunostaining specifically target to the antigen/protein of interests by our previous studies (PMID:26839042; PMID: 27143681; PMID: 29664678; PMID: 29924941). The anti-NG2 antibodies were well validated by immunostaining, western blotting and flow cytometry (PMID: 28964848; PMID: 31131431; PMID:31776510) and was not colocalized with other cells including arterial ECs, smooth muscle cells based on our studies. The anti-TMEM100 antibodies were previously validated by immunostaining on Tmem100 CKO mice (PMID: 25640077) showed Tmem100 CKO mice did not express Tmem100. Our data showed that TMEM100 is enriched in human lung but not expressed the kidney and liver ECs. The immunostaining showed similar data suggest that the TMEM100 antibodies specifically label TMEM100.

**RNA-Seq analysis**

HPMVECs with TMEM100 or Vector overexpression were lysed for RNA isolation with the Zymo Research Quick-RNA Miniprep including DNase I digestion. Equal amounts of RNA from ECs isolated from three individual GFP or TMEM100 overexpression were pooled and sequenced with NovaSeq PE150 at Novogene Corporation Inc. (Sacramento, CA, USA) The original sequencing data were aligned with HISAT2[5], and the differential expression analysis was performed using Cuffdiff software[6]. Kyoto Encyclopedia of Genes and Genomes (KEGG) pathways analysis were analyzed based on the upregulated genes [Fragments Per Kilobase of transcript per Million mapped reads (FPKM) > 1 & fold change > 1.5] using DAVID Bioinformatics Resources (https://david.ncifcrf.gov).

**QRT-PCR analysis**

QRT-PCR was performed on QuantStudio 3 Real-Time PCR System (Thermo Fisher Scientific, Waltham, MA, USA) with PowerUp SYBR Green Master Mix (Thermo Fisher Scientific, Waltham, MA, USA). Target mRNA was determined using the comparative cycle threshold method of relative quantitation. 18s rRNA gene was used for human genes. The primer sequences are provided in Supplemental Table 2.

**BrdU assay**

HPMVECs were infected with Vector or TMEM100 lentiviruses for 48 h. 5-bromo-20-deoxyuridine (BrdU, 10 μM, Sigma) was added for the 4 h. Cells were stained with anti-BrdU (BD Biosciences, Cat#BDB347580, 5 $\mu$g/ml) according to the manufacturer's instructions and nuclei were counterstained with DAPI. Images were taken using REVOLVE microscopy (Discover Echo Inc.) at 4 X magnification for quantification.

**Tube formation assay**

HPMVECs were infected with Vector or TMEM100 lentiviruses for 48 h. Cells were detached and seeded on Matrigel (Sigma, Cat# E1270-5ML) at the density of 40,000 cells per 96 well. Two to 4 hours later, Images were taken using REVOLVE microscopy (Discover Echo Inc.) at 10 X magnification for quantification of branch points.

**Transendothelial Monolayer Electrical Resistance (TER) Assay**

Endothelial monolayer junctional changes were measured by an electric cell substrate impedance sensor (ECIS) to record real-time changes in electrical resistance across endothelial monolayers. Briefly, HPMVECs after infection of Vector and TMEM100 lentiviruses for 48 hours were seeded at confluence on a ECIS chamber with small gold electrode (Applied Biophysics) overnight. Before the experiment, the confluent endothelial monolayer was monitored for 0.5 hour at baseline, the endothelial monolayers were treated with Thrombin (4U/ml) and monitored up to 12 hours. The data was normalized to the baseline value.

**Western Blot**

Mouse lung tissues were well perfused with 1X PBS and collected for homogenization in NP-40 buffer supplemented with protease inhibitor cocktails (Sigma, Cat#5391371ML). Western blotting assays were performed using anti-TMEM100 (Millipore Sigma, Cat#ABN1721, 1 $\mu$g/ml), anti-VE-cadherin (Thermo Fisher, Cat# 361900, 0.5 $\mu$g/ml), anti-Flag (Sigma, Cat#F1804, 0.5 $\mu$g/ml ), Anti-β-actin (Sigma, Cat #A2228, 0.2 $\mu$g/ml) or anti-GAPDH (Santa Cruz, Cat# SC-47724, 0.04 $\mu$g/ml) was used as a loading control.

**Statistical Analysis**

Statistical determination was performed on Prism 9 (Graphpad Software Inc.). Two-group comparisons were compared by parametric test with the unpaired 2-tailed Student t test for equal variance and Welch's t-test for unequal variance. Based on additional literature support from similar studies, our samples fit normal distribution. P less than 0.05 indicated a statistically significant difference. All bars in dot plot figures represent mean. All bar graphs represent mean±SD. In DAVID pathway analysis, Fisher's Exact test is adopted to measure the gene-enrichment in annotation terms.

**Sample size estimation, inclusion/exclusion criteria, and randomization**

Sample size estimation and randomization were not performed. Sample size was determined based on the literature and our previous experience. All the samples performed were included. No data was excluded.

### Online Figure and legend

**Supplemental Table 1, human sample information**

| Organ | Age (year) | Sex | Sample source |
|---|---|---|---|
| Lung | 13 | Male | PHBI |
| Lung | 21 | Male | PHBI |
| Lung | 10 | Male | CCMC |
| Lung | 15 | Male | CCMC |
| Brain | 15 | Male | CCHC |
| Kidney | 16 | Female | CCHC |
| Heart | 16 | Female | CCHC |
| Liver | 16 | Female | CCHC |

**Supplemental Table 2, Primer sequences for QRT-PCR analysis**

| Gene name | Forward primer | Reverse primer |
|---|---|---|
| *hFOXM1* | GGAGGAAATGCCACACTTAGCG | TAGGACTTCTTGGGTCTTGGGGTG |
| *hE2F1* | ACTGACCATCAGTACCTGGC | CGGGGATTTCACACCTTTTCC |
| *hPLK1* | CACAGTGTCAATGCCTCCAAG | TATCACAGAGCTGATACCCAAG |
| *hCDKN2C* | ACTGCGCTGCAGGTTATGAA | AGCGAAACCAGTTCGGTCTT |
| *hCCNB1* | ACCTGTGTCAGGCTTTCTCTG | CTGACTGCTTGCTCTTCCTCA |
| *hCCNB2* | GCACAAGTAGCTAAGAAAGCTCA | CTCAGGTGTGGGAGAAGGAC |
| *hCCNA2* | CTCAGAAGAAGCCAGCTGAATC | CTGTTAGTGATGTCTGGCTGT |
| *h18S rRNA* | TTCCGACCATAAACGATGCCGA | GACTTTGGTTTCCCGGAAGCTG |

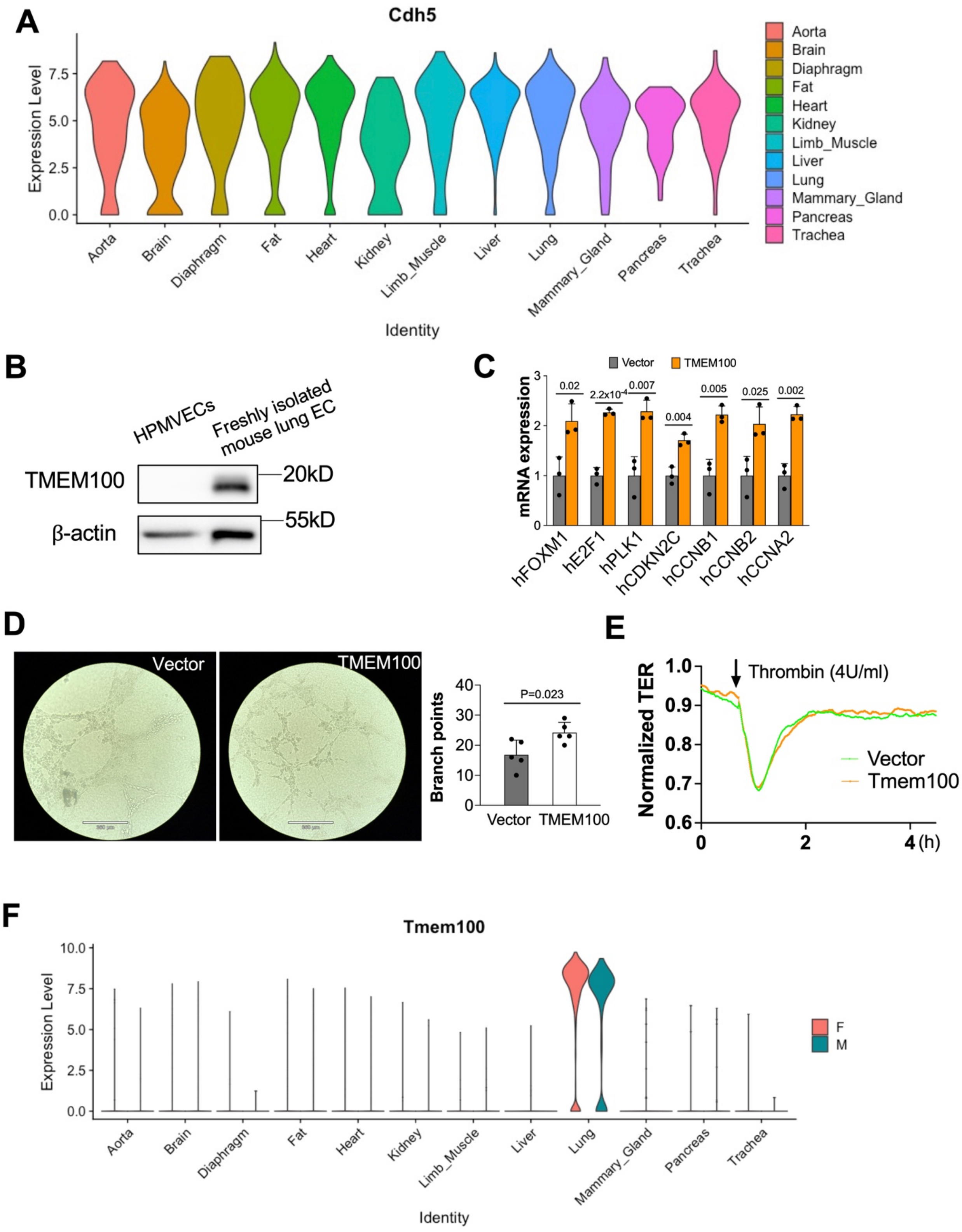
A
Cdh5
Expression Level
Identity
Aorta
Brain
Diaphragm
Fat
Heart
Kidney
Limb_Muscle
Liver
Lung
Mammary_Gland
Pancreas
Trachea
B
HPMVECs
Freshly isolated mouse lung EC
TMEM100
20kD
β-actin
55kD
C
Vector
TMEM100
mRNA expression
0.02
2.2x10-4
0.007
0.004
0.005
0.025
0.002
hFOXM1
hE2F1
hPLK1
hCDKN2C
hCCNB1
hCCNB2
hCCNA2
D
Vector
TMEM100
Branch points
P=0.023
E
Thrombin (4U/ml)
Normalized TER
Vector
Tmem100
(h)
F
Tmem100
Expression Level
F
M
Identity

**Supplemental Figure A. ScRNA-seq analysis showing the expression of pan EC marker CDH5 across different mouse organs.** Tabula Muris EC scRNA-seq dataset was used for analysis.

**B. TMEM100 is not expressed in the cultured HPMVECs but highly expressed in the isolated mouse lung ECs.** HPMVECs were cultured in vitro at passage 8. Lung ECs were isolated via CD31 conjugated microbeads.

**C. QRT-PCR analysis demonstrated that TMEM100 overexpression upregulated genes related to cell proliferation.** HPMVECs were infected with TMEM100 lentiviruses for 48 hours. Cells were starved overnight and used for QRT-PCR analysis. N=3.

**D. Tube formation assay showed that TMEM100 overexpression induced angiogenesis in vitro.** HPMVECs were infected with TMEM100 lentiviruses for 48 hours. Cells were transferred to Matrigel for tube formation assay. N=5. Student t test.

**E. Transendothelial Monolayer Electrical Resistance (TER) Assay demonstrated that TMEM100 overexpress did not affect cellular junction.** TMEM100 overexpression did not affect cell junction integrity in HPMVEs. At 48 hours post-transfection, TER was monitored for up to 5 hours. Thrombin (4U/ml) was added to disrupt the cellular junction. N=3.

**F. There is not difference in Tmem100 expression between male and female mice.** Tabula Muris EC scRNA-seq dataset was used for analysis.
